# Lung epithelial progenitor-mediated release of TGF-β regulates induction and persistence of lung CD8^+^ T_RM_ cells following mucosal BCG vaccination

**DOI:** 10.1101/2024.10.16.618714

**Authors:** Judith A. Blake, Julia Seifert, Roland Ruscher, Paul R. Giacomin, Denise L. Doolan, Andreas Kupz

## Abstract

A principal reason for the high global morbidity and mortality of tuberculosis (TB) is the lack of efficacy of the only licensed TB vaccine, Bacillus Calmette-Guérin (BCG), as parenteral BCG does not induce local pulmonary immune memory. Animal studies have shown that mucosal BCG vaccination provides superior protection against TB due to generation of lung resident memory T cells (T_RM_). Here, we demonstrated that following mucosal vaccination with the genetically modified virulent BCG strain, BCG::RD1, distal airway epithelial progenitors were mobilized to assist with restoration of alveolar epithelium. By way of their migration-mediated activation of latent TGF-β, lung CD8^+^ T_RM_ differentiation was induced. Mucosal vaccinations using nonvirulent strains of BCG in which airway epithelial progenitors were not mobilized, as well as genetic inhibition of migration-mediated activation of TGF-β, resulted in significantly lower numbers of lung CD8^+^ T_RM_. In addition, we discovered CD8^+^ cells with ex-lung and stem-like T_RM_ phenotypes that persisted in the lung-draining mediastinal lymph nodes for up to four months following mucosal BCG vaccination. These results link airway epithelial progenitor-mediated repair of injured lung tissue with induction of resident T cell memory and delineate why persistence of T_RM_ in the lung is short-lived. These findings may explain why mucosal vaccination with virulent BCG strains is more protective against TB and thus have notable implications for future TB vaccine development.

**One Sentence Summary:** Following lung damage due to inhalation of virulent BCG, distal airway epithelial progenitor cells interact with lung CD8^+^ T cells to induce their differentiation into resident memory T cells via migration-mediated activation of TGF-β.

## INTRODUCTION

During the Covid-19 pandemic, the ineffectiveness of a parenteral vaccine to protect against lung infection became more apparent, a problem that was well acknowledged amongst TB immunologists (*1*). TB is primarily a respiratory infection caused by *Mycobacterium tuberculosis* (*Mtb*); boasting 10.6 million new infections in 2022 and 1.3 million deaths, TB is one of the top ten fatal diseases globally (*2*). These statistics highlight the inadequacy of the only licensed TB vaccine - BCG (*1, 3, 4*). BCG is an attenuated strain of *Mycobacterium bovis* given parenterally to postnatal infants in TB endemic regions to confer immunity against acute septic forms of TB (*5*), however, immunity is lost in adolescence (*6*). Failure of BCG to provide life-long immunity against *Mtb* is thought due to insufficient generation of antigen-specific CD8^+^ memory T cells in the lungs (*7*). This is an area of vaccine improvement where recombinant modifications of BCG aim to generate a more persistent CD8^+^ T cell memory. For example, mice and guinea pigs vaccinated with BCG strains in which the region of difference 1 genes that encode *Mtb* virulence genes ESAT-6 and CFP-10 were recombined into *M. bovis* BCG (BCG::RD1) (*8*) showed improved protection against *Mtb* challenge (*9, 10*).

A major goal of vaccination is to induce antigen-experienced circulatory and tissue-resident memory T cells (T_RM_) that respond rapidly to secondary infection. It has been demonstrated that a mucosal (or inhaled) BCG vaccination, as opposed to parenteral BCG, generated superior protection against TB in animal models and this protection was linked to generation of lung T_RM_ (parenteral BCG did not lead to T_RM_ in the lungs) (*9, 11*). Mucosal delivery of BCG::RD1 has been further shown to induce significant numbers of airway T_RM_ (*12*) and conferred superior immunity against aerosol *Mtb* infection in diabetic mice (*13*). Thus, immunogenic antigens of BCG::RD1 (*9*) combined with a mucosal delivery strongly contributes to vaccine-induced protection against *Mtb*.

Stem/progenitor cells perceive damage, and designate resident and recruited immune cell roles. Evidence of interaction between immune and progenitor cells is seen in the array of immune receptors and cytokine signalling systems of progenitor cells (*14–16*) and dysregulation of immune-progenitor cell interaction results in chronic disease (*17–19*). In mouse models of acute influenza infection, there is an invariable association of CD8^+^ T_RM_ and lung epithelial progenitor cells in areas of severe tissue destruction (*20–27*). In that context it has been suggested that “auxiliary developed regenerative tissues could be a source of unique factors necessary for differentiation and maintenance of T_RM_ cells, such as TGF-β” (*28*).

Lung injury activates adult epithelial progenitors to regenerate the alveolar epithelium which consists of Type 1 and Type 2 alveolar epithelial cells (AEC1 and AEC2, respectively). AEC1 are responsible for gas exchange while AEC2 secrete surfactant to prevent alveolar collapse, amongst many other functions. Using a Cre-reporter for Wingless-related integration site (Wnt) signalling, a rare AEC progenitor (AEP) that constitutes ∼20% of all AEC2 was described (*29*). Although considered stem cells, AEP express AEC2 markers and function as AEC2 (*23*). Zacharias *et al*. (*23*) reported the distinct functional AEP gene expression profile enriched in lung developmental genes and epigenome. Thus, a small subset of AEC2 termed AEP possess bipotency that are recruited to repair alveolar damage.

Following severe injury and depletion of normally abundant local alveolar epithelial progenitor cells, wound-healing responses originate from stem cells in the airways (*30*). Of relevance to our studies are lineage negative epithelial progenitors (LNEP aka Sox2^+^ Lin^-^) and distal airway stem cells (DASC). LNEP in the distal bronchioles are quiescent in uninjured lungs, however, alveolar injury can elicit their proliferation, such that daughter cells migrate into alveolar tissue and differentiate into AEC1 or AEC2 (*31, 32*). LNEP are targeted by murine PR8 influenza virus and lose their proliferative capacity (*33*); therefore, rare, quiescent DASC proliferate, and daughters migrate to occupy alveolar areas depleted of epithelium where they form Krt5^+^ “pods” with CD8^+^ T_RM_ (*22, 34*).

How T_RM_ induction and maintenance is regulated following mucosal exposure to mycobacteria, particularly parental and recombinant strains of BCG, and if it is linked to airway progenitor-mediated restitution of injured lung tissue, remained unknown. Thus, we studied CD8^+^ lung T_RM_ and lung epithelial progenitors to elucidate how they cooperate for restoration of lung function following inhaled BCG vaccination. We describe a novel mechanistic association between airway epithelial progenitor paracrine cues and induction of CD8^+^ lung T_RM._

## RESULTS

### BCG virulence and dose affects loss of alveolar epithelial cells

To study the response of lung epithelial cells to mucosal vaccination with BCG strains across a spectrum of dose and virulence over time, we used intratracheal (IT) vaccination of C57BL/6 mice (n=6/time point) with the Danish strain of BCG (BCG SSI) at a dose of 100,000 colony forming units (CFU), BCG SSI at 500,000 CFU, BCG::RD1 at 100,000 CFU or PBS (Fig. 1a).

**Figure 1.**
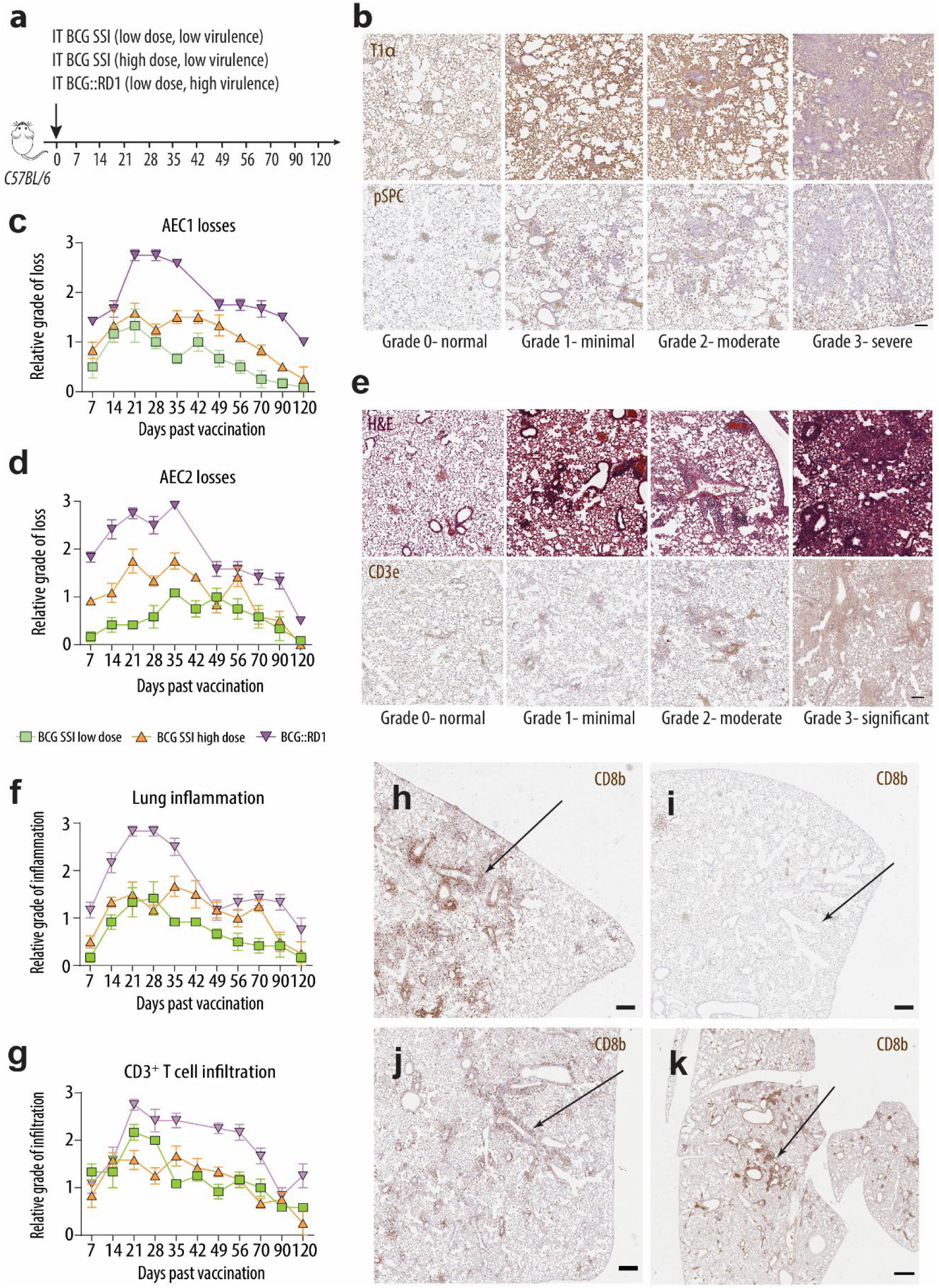
AEC losses, inflammation, and acquired immune responses are directly related to IT BCG dose, virulence, and delivery. **a)** Experimental design to define AEC losses and immune responses. **b)** Examples of semi-quantitative tissue grading of AEC1 and AEC2 losses using immunohistochemical (IHC) staining for T1α and pSPC, respectively. Semi-quantitative enumeration of **c)** AEC1 and **d)** AEC2 losses over time. **e)** Examples of semi-quantitative alveolar tissue grading for inflammation and infiltration of T cells using H&E and CD3e, respectively and **f, g**) semi-quantitative data. Representative images of distal alveolar epithelium and terminal bronchioles (arrows) show significantly more infiltration by CD8^+^ T cells in the **h)** BCG::RD1 and **j)** BCG SSI low dose vaccinated lungs as compared to **i)** BCG SSI high dose. **k)** Proximal sections of lungs vaccinated with BCG SSI high dose show CD8^+^ T cell infiltration. Results are presented from representative images and pooled data means ± SEM from 3 mice per time group from two pooled independent experiments (n=6). **b)** Magnification, 5.2x; Scale bar, 400μm. **e)** Magnification, 4.2x; Scale bar 500µm. **h-j)** Magnification, 2.5x; Scale bar 900µm. **k)** Magnification, 1.5x; Scale bar, 2mm.

Mycobacteria infect AEC2 following inhalation, so AEC2 are frontline cells of invasion (*35*); also, loss of cell-to-cell contacts due to infection-related cytolysis of AEC2 also affects AEC1 that are particularly sensitive to epithelial damage (*36*). To determine the relative number of AEC lost (the impetus for immune and subsequent progenitor cell responses) across our vaccinations, lung sections were immunostained using type 1 alpha protein (T1α) and pro-surfactant protein C (pSPC), before semi-quantitative grading of AEC1 and AEC2 losses, respectively (Fig. 1b).

Following BCG SSI low dose vaccination, minimal AEC1 losses were observed by the peak of injury at 14 days past vaccination (dpv) (Fig. 1c). Similarly, AEC2 loss was minimal by the peak at 28-35 dpv (Fig. 1d). AEC pathology consisted of sparse small focal lesions (Supp. Fig. 1a), as well as alveolar wall hyperplasia where alveoli structurally stayed intact. Following BCG SSI high dose vaccination, moderate AEC1 loss was observed by the peak of injury at 21 dpv (Fig. 1c). Moderate AEC2 losses appeared from 21 dpv through to 70 dpv before clearing at 120 dpv (Fig. 1d). AEC pathology consisted of global focal peribronchiolar lesions (Supp. Fig. 1b). After BCG::RD1 vaccination, a marked AEC1 loss occurred by a peak at 21 dpv, before reduction to moderate losses at 49 dpv and continual minimal losses until 120 dpv (Fig. 1c). Marked widespread loss of AEC2 was observed from 21 to 35 dpv and remained moderate from 49 dpv to 70 dpv with further minimal losses until 120 dpv (Fig. 1d). AEC pathology consisted of large global areas of severe destruction of both AEC with necrosis evidenced by T1α^+^ and pSPC^+^ cell debris in bronchiolar lumen (Supp. Fig. 1c). Taken together, these results illustrated a direct relationship between dose and virulence of the BCG vaccine and AEC losses.

### BCG dose, virulence and delivery impacts lung inflammation and T cell influx

To broadly define immune responses to AEC losses, lung sections from each animal were stained with hematoxylin and eosin (H&E) and immunostained with CD3 epsilon (CD3e), before semi-quantitative grading of inflammation and T cell infiltration, respectively (Fig. 1e). After BCG SSI low dose vaccination, inflammation rose slowly to peak at moderate levels by 28 dpv and largely resolved by 35 dpv (Fig. 1f). T cells infiltrated relatively early compared to the other two vaccines, reaching moderate numbers by 21 dpv and declining steadily thereafter (Fig. 1g). The inflammatory response consisted of small haemorrhagic lesions (Supp. Fig. 1d) due to vascular changes and minimal focal peribronchiolar and perivascular inflammation. When BCG SSI was administered at a high dose, inflammation and T cell infiltration did not peak until 35 dpv (Fig. 1f, g). The inflammatory response consisted of global focal areas of haemorrhage due to capillary changes and moderate peribronchiolar (primarily T cells) and perivascular immune cell infiltrates, as well as moderate numbers of interstitial giant cell macrophages (Supp. Fig. 1e). Following BCG::RD1 vaccination, global inflammation escalated to severe levels by 21 dpv and remained severe until 42 dpv (Fig. 1f). T cell infiltration also peaked by 21 dpv and T cells continued to infiltrate in high numbers until 56 dpv (Fig. 1g). The inflammatory response consisted of haemorrhage secondary to inflammation, marked peribronchiolar and perivascular immune cell infiltrates, and marked numbers of interstitial giant cells and foamy macrophages.

We observed that at the terminal bronchioles, the BCG SSI high dose vaccine did not induce infiltrating CD8^+^ T cells (Fig. 1i). This was in marked contrast to that seen with the BCG::RD1 (Fig. 1h) and low dose BCG SSI (Fig. 1j) vaccinations. We reasoned that numbers of bacilli made the vaccine more viscous and therefore it was not delivered into the distal airways; indeed, lungs sectioned up to proximal airways showed infiltration of T cells in peribronchiolar regions (Fig. 1k). With these experiments to establish the timeline of immune responses, we demonstrated that dose, virulence, and delivery of mucosal BCG vaccination had direct effects on inflammation and acquired immune responses.

### Only Sox2^+^ distal airway and alveolar epithelial progenitors respond to AEC losses associated with mucosal BCG vaccination

We next undertook an original characterization of lung epithelial progenitor cell responses to mucosal BCG vaccinations. Lung sections were probed using antibodies against Krt5 for DASC, Sox2 for distal airway epithelial progenitors and pSPC for alveolar progenitors across the timeline. Results demonstrated that although Krt5 was expressed in basal cells of the large conducting airways, we did not observe Krt5^+^ cells in alveolar tissue at any time after vaccination (Fig. 2a). We confirmed these findings using *Krt5^CreERT2^/mTmG* reporter mice to indelibly tag DASC progeny with green fluorescent protein (*37*). Again, we observed green fluorescent protein in the large airway basal cells but never in alveolar tissue (Fig. 2b).

**Figure 2.**
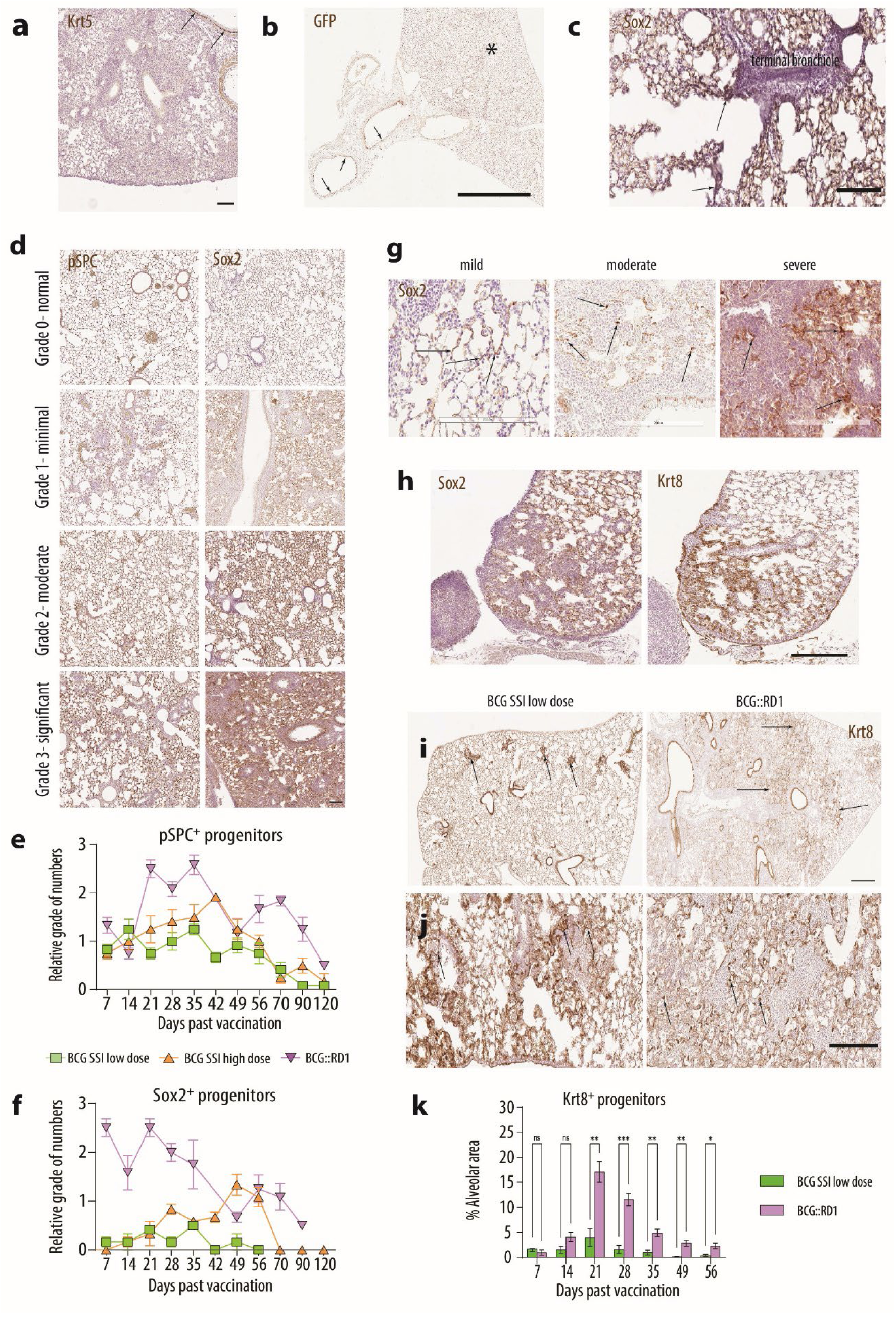
Activation of distal airway and alveolar progenitors is dependent upon dose and virulence of IT BCG vaccination. **a)** Krt5 and **b)** GFP immunoreactivity in the large conducting airways (arrows) was not seen in alveolar regions (asterisk) at any timepoint following any IT BCG vaccination. **c)** Representative image at 7 days after BCG::RD1 vaccination, where Sox2 expression is seen in cells (arrows) in and adjacent to terminal bronchioles **d)** Examples of semi-quantitative alveolar tissue grading for proliferation of alveolar and distal airway progenitors using pSPC and Sox2 IHC staining, respectively. **e, f)** Semi-quantitative data are presented from representative images and pooled data means ± SEM from 6 mice per time group from two independent experiments. **g)** Sox2^+^ airway progenitors (arrows) were observed throughout all areas of alveolar damage. Krt8 staining revealed that **h)** the TP state was observed within tissue areas of Sox2 expression, in an ‘inverse’ relationship. **i)** Lungs 14 days after vaccination with BCG SSI showed punctate areas of Krt8^+^ TP (arrows), while those from BCG::RD1 lungs showed diffuse Krt8^+^ TP throughout (arrows). **j)** 21 days after BCG SSI vaccination, Krt8^+^ cells had a squamous morphology (arrows) while in BCG::RD1 vaccinated lungs, Krt8^+^ TP retained round morphology (arrows). **k)** Quantification of Krt8^+^ TP numbers. Results are presented from representative images and data means ± SEM from 3 mice per group from two pooled independent experiments (n=6). \**P* < 0.05; \*\**P* < 0.01; \*\*\**P* <0.001; ns: not significant by unpaired Student’s *t* test. **a, c, d, h, j)** Magnification, 10x; Scale bar 200µm. **b)** Magnification, 1.5x; Scale bar 2mm. **g)** Magnification, 20x; Scale bar 200µm. **i)** Magnification, 4.2x; Scale bar 500µm.

In contrast, by 7 dpv after BCG::RD1 vaccination, rounded cells with nuclear Sox2^+^ expression were observed in terminal bronchioles; in contiguous alveolar epithelium, squamous cells with cytoplasmic Sox2^+^ expression were also noted (Fig. 2c). Therefore, across our vaccination’ timeline, relative enumeration of Sox2^+^ distal airway and pSPC^+^ alveolar epithelial progenitors were undertaken using the respective markers and semi-quantitative grading for relative numbers of the two progenitor cell types (Fig. 2d).

Results demonstrated that following the BCG::RD1 vaccination significant numbers of Sox2^+^ airway progenitors were observed early (Fig. 2f) but subsequently waned after day 21; conversely, numerous pSPC^+^ alveolar progenitors were observed only after day 21 (Fig. 2e). In areas of severe destruction, pSPC^+^ progenitors were not seen, although they were on the periphery of parenchymal lesions. Of note, Sox2^+^ cells were seen throughout all areas of alveolar damage (Fig. 2g). Following BCG SSI high dose vaccination, Sox2^+^ airway progenitors peaked later at 49 dpv (Fig. 2c) probably related to the later peak of cell losses, inflammation, and T cell responses; these cells were confined largely to proximal peribronchiolar regions. After BCG SSI low dose vaccination, pSPC^+^ alveolar epithelial progenitors appeared largely as doublets with ‘foci’ of cells across neighbouring alveoli as described by Desai *et al.* (*38*); Sox2^+^ distal airway progenitor numbers were minimal (Fig. 2c).

More recently, a Krt8^+^ transitional progenitor (TP) cell state that precedes differentiation of both Sox2^+^ airway and pSPC^+^ progenitors into AEC1 was described (*32, 39, 40*). We investigated Krt8^+^ TP using our vaccinations as we reasoned that these epithelial progenitors interact with lung CD8^+^ T_RM_. In serial sections, we observed an inverse staining pattern in which alveolar areas void of Sox2 staining had Krt8 staining and vice versa (Fig. 2h). This supported the notion that Sox2^+^ airway cells were transitioning into Krt8^+^ progenitors. By 14 dpv, the BCG SSI low dose had punctate areas of Krt8^+^ TP, while the BCG::RD1 vaccination had broad diffusion of Krt8^+^ TP throughout lungs (Fig. 2i). At 21 dpv after BCG SSI low dose, Krt8^+^ cells acquired a squamous morphology and were likely transitioning into AEC1 (Fig. 2j); conversely, after BCG::RD1 vaccination, Krt8^+^ cells remained round (Fig. 2j) and did not become squamous until after 28 dpv (data not shown). Quantification of Krt8^+^ TP for both vaccinations showed a peak of Krt8^+^ TP at 21 dpv where BCG::RD1 vaccinated lungs had an average alveolar area of ∼20% occupied by Krt8^+^ cells, while those of the SSI low dose had an area of ∼5% (Fig. 2k).

In summary, responses of AEP, LNEP, and DASC to alveolar epithelial cell loss associated with mycobacterial lung injury were unknown at the start of this research; moreover, the literature did not describe colocalization of CD8^+^ T_RM_ and epithelial progenitor cells following mycobacterial lung infection. Knowing the effects that RD1 virulence factors have on AEC2, we initially presumed that vaccination with BCG::RD1 may cause a similar lung injury healing sequalae as reported after murine virulent influenza (PR8) infection in mice (*20, 22, 23, 33, 41*). Our findings, instead, described sequalae in which Sox2^+^ distal airway progenitors (we suggest the progeny of LNEP) migrated into alveolar tissues to orchestrate restitution in areas where facultative alveolar epithelial progenitors were lost due to the virulence factors of RD1. Conversely, mucosal lung exposure to doses of non-virulent BCG resulted in efficient regeneration of lost epithelium by local epithelial progenitors.

Thus, we demonstrated that solely Sox2^+^ distal airway and alveolar progenitors responded to AEC losses associated with mucosal BCG vaccination. Where alveolar tissue damage was mild or moderate, alveolar epithelial progenitors were observed, whereas, where alveolar tissue damage was severe, Sox2^+^ distal airway progenitors were observed. Both progenitor types likely transitioned through the Krt8^+^ TP state into AEC1, evidenced by near complete tissue restitution at 120dpv.

### Lung CD8^+^ T_RM_ induction depends on BCG virulence and dose

The pulmonary architecture is thought to be non-conducive to a permanent anatomical niche for CD8^+^ T_RM_ to function, such as that known for skin. It is intuitive that over the lifespan of a human whose lungs encounter a myriad of respiratory pathogens, numerous depots of accumulative parenchymal CD8^+^ T_RM_ could be deleterious for the delicate architecture and surface area required for efficient gas exchange. Undeniably, lung CD8^+^ T_RM_ have unique organ-specific mechanisms for maturation, persistence, and egress by which ‘the complexity of this system and interdependence between the various components and changes that affect parameters such as the timing of T-box TF modulation and the coordination with TGF-β availability would have a profound effect on T_RM_ cell maturation’(*42*). Thus, our next aim was to define induction and persistence of CD8^+^ T_RM_ subsets following vaccination with the three different BCG dose/virulence combinations. Mice were given IT BCG vaccination as described (Fig. 1a); whole lungs of experimental animals were harvested and processed into single-cell suspensions before CD8^+^ T_RM_ quantification using flow cytometry (Fig. 3a).

**Figure 3.**
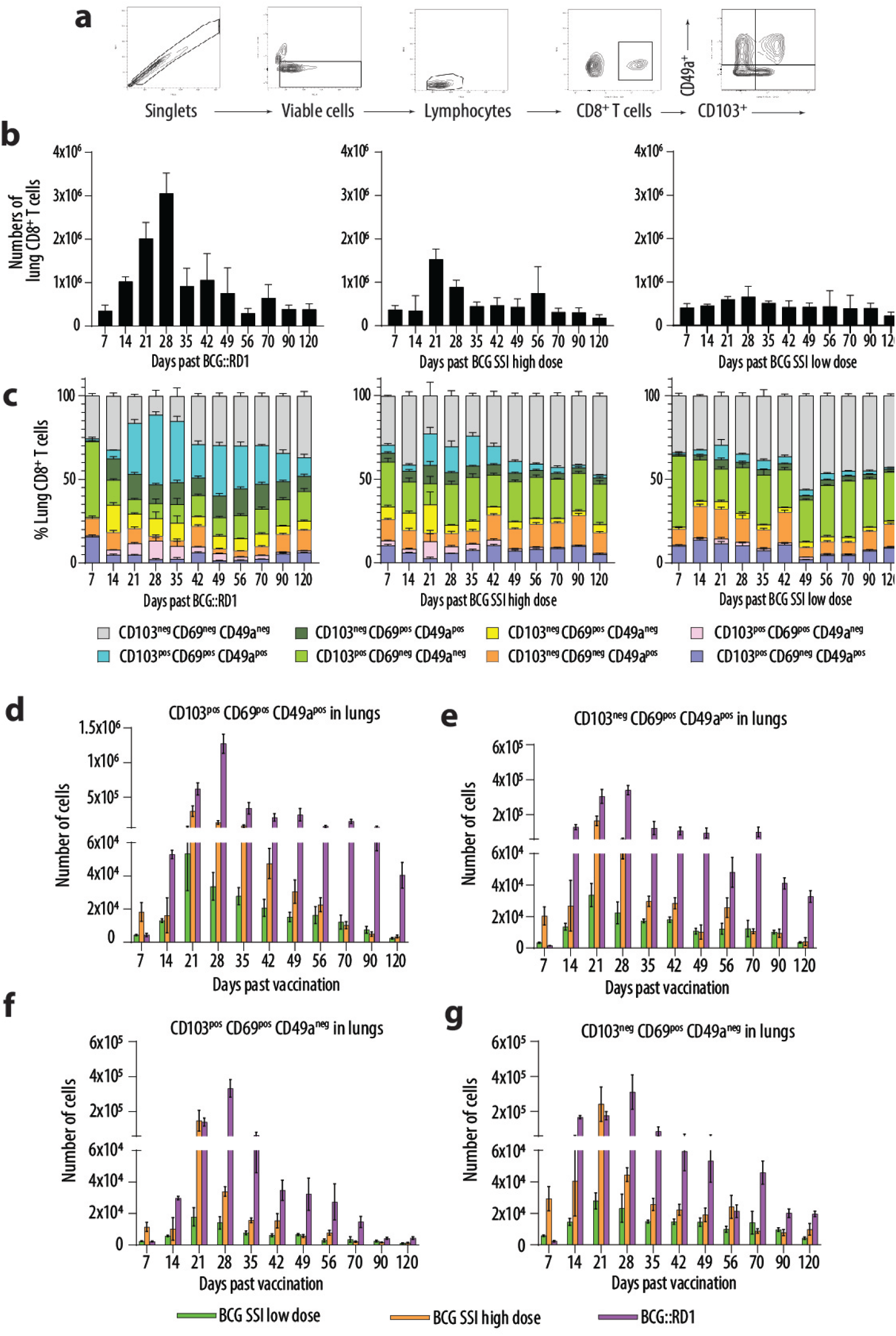
CD8α^+^ T cell numbers and percentages of lung CD103, CD69 and CD49a T_RM_ subsets across models of mucosal BCG vaccination. **a)** Flow cytometry gating strategy for CD8^+^ T_RM_ quantification. Flow cytometry analysis of **b)** numbers of CD8α^+^ T cells; **c)** the percentages of CD8α^+^ T_RM_ subsets and **d-g)** numbers of CD8^+^ T_RM_ subsets in the lungs of vaccinated mice. Pooled data from two independent experiments where n=4-10 for each timepoint.

BCG::RD1 vaccination led to robust expansion of CD8^+^ T_RM_ from days 14-35 dpv, with a peak at day 28; numbers contracted at day 35 yet remained high until 120 days (Figs. 3d-g). Of the three subsets of T_RM_, CD103^+^CD69^+^CD49a^+^ T_RM_ cells that patrol airway/ interstitial borders (*43*) were the highest proportion at 45% of total CD8^+^ T cells at their peak at 28 dpv (Fig. 3b-d). Interstitial CD103^-^CD69^+^CD49a^+^ T_RM_ (*43, 44*) were also abundant (Fig. 3c, e) while the short-lived CD103^+^CD69^+^CD49a^-^ T_RM_ that solely patrol the airways (*43, 45*) were considerably less (Fig. 3c, f). CD103^-^CD69^+^CD49a^-^ T_RM_ precursor cells (*44*) were continually present in the lung from day 14-120 (Fig. 3c, g).

After vaccination with a high dose of BCG SSI, CD8^+^ T cells in the lung were half the numbers seen following BCG::RD1 vaccination (Fig. 3b). Of those, 25% were CD103^+^CD69^+^CD49a^+^ T_RM_ at day 21 dpv (Fig. 3b-d) while CD103^+^CD69^+^CD49a^-^ airway effector T_RM_ were higher than those seen after BCG::RD1 vaccination (Fig. 3c, f). BCG SSI low dose vaccination induced only about a third of CD8^+^ T cells in the lung compared to BCG::RD1 vaccination (Fig. 3b, c). Of those, 10% were CD103^+^CD69^+^CD49a^+^ T_RM_ at day 21 dpv (Fig. 3c, d) and waned each week. Taken together, we demonstrated that lung CD8^+^ T_RM_ induction was virulence- and dose-related in all vaccination regimens and that CD8^+^ T_RM_ waned in the lungs by 120 days after mucosal BCG vaccination.

### Lung-draining mediastinal lymph node harbor CD8^+^ T_RM_ following mucosal BCG vaccination

It has been consistently demonstrated that lung CD8^+^ T_RM_ wane over a short time (*11, 46–48*), despite continual circulation of antigen-specific T cells in blood (*21*). It has also been suggested that waning of local immune memory in lung is an organ-intrinsic protective mechanism which safeguards against the potential of CD8^+^ T_RM_ in successive infections to mediate considerable tissue damage (*49, 50*) as was seen in persons with successive SARS-CoV-2 infections (*51, 52*). As lung CD8^+^ T_RM_ are known to egress to the lung-draining mediastinal lymph node (mLN), where they are thought to provide regional immune memory for the lungs (*50*), we next investigated whether this was the case following mucosal BCG vaccination. Mice were given IT vaccination as described, the right superior mLN was extracted from all experimental animals across 7-120 dpv and processed into single-cell suspensions before quantification of CD8^+^ T_RM_ using flow cytometry.

We discovered all phenotypes of ex-lung CD8^+^ T_RM_ in the mLN. CD103^+^CD69^+^CD49a^+^ and CD103^+^CD69^+^CD49a^-^ T_RM_ were conceivably swept into lymphatics due to tissue damage that physically dislodged them (*49*) (Fig. 4a) supported by the observation of these subsets early, associated with the time of severe damage following BCG::RD1 vaccination (Fig. 4b, d) and later after BCG SSI high dose vaccination, likely associated with its characteristic late immune response (Fig. 4b, d).

**Figure 4.**
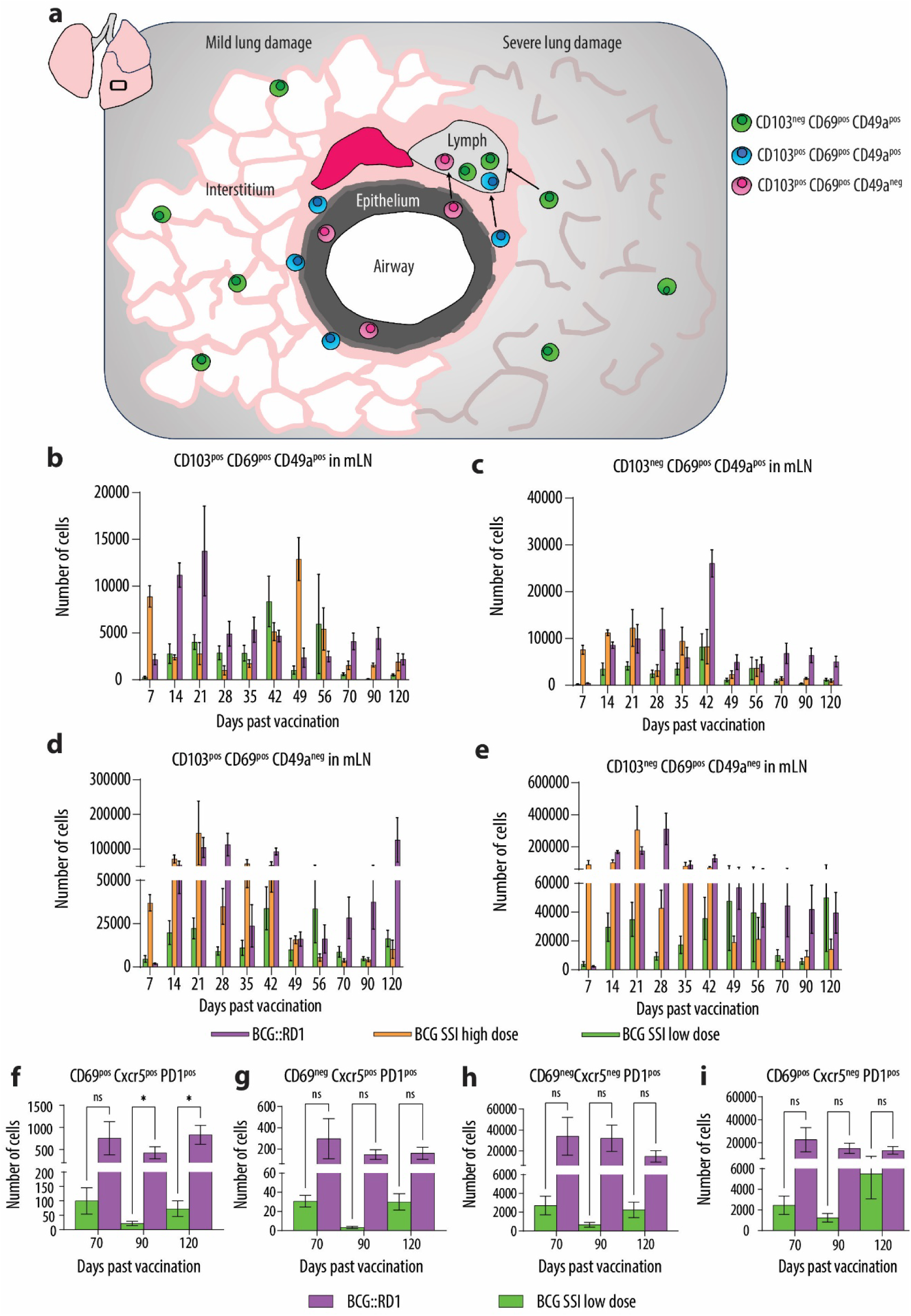
Ex-lung and stem-like CD8^+^ T_RM_ counts in mLN after mucosal BCG vaccinations. **a)** Diagram of distribution of CD8^+^ T_RM_ subsets in mildly damaged lungs and their dissociation into lymphatics in severely damaged lungs. **b-e)** Flow cytometry quantification of CD8^+^ T_RM_ subsets in mLN of vaccinated mice. Flow cytometry quantification of mLN **f)** quiescent, **g)** proliferative stem-like CD8^+^ T_RM_ and **h, i)** reinvigorated effector T cell progeny phenotypes. Pooled data of mLN CD8^+^ T_RM_ from two independent experiments where n=4-10 for each timepoint. \**P* < 0.05; ns: not significant by unpaired Student’s *t* test.

Interstitial CD103^-^CD69^+^CD49a^+^ T_RM_ (Fig. 4a) lack CD103 maturation which is said to contribute to their egression to the mLN (*49*). In addition, when restimulated, CD103^+^CD69^+^CD49a^+^ T_RM_ proliferate within the mLN and lose CD103 expression (*53*). Here, we showed for the first time that following mucosal BCG vaccination, mLN harbor these CD103^-^CD69^+^CD49a^+^ T_RM_ which, importantly, may have the capacity to repopulate the lungs as T_RM_ following secondary infection (*54–56*) (Fig. 4c); these will be a focus of our future studies.

We also noted CD103^-^CD69^+^CD49a^-^ of the ‘precursor’ T_RM_ phenotype (*44*) at the latest stages of the timeline in the mLN (Fig. 4e) when numbers of lung CD8^+^ T_RM_ had steadily declined, and tissue was largely restituted. We reasoned that these may be exhausted effector T cells that home to B cell follicles of draining lymph nodes due to chronic antigen stimulation where they become stem-like CD8^+^ T_RM_ (*57–59*). With chronic *Mtb* infection, prolonged overexposure to antigens has been reported to cause T cells to lose effector functions and exhibit an exhausted phenotype (*60*) including expression of PD-1 (*61*); also, anti-TB therapy has been shown to promote reduced expression of PD-1 and its ligands (*62, 63*).

As previous studies have shown the persistence of viable BCG in IT vaccinated lungs of mice up for to 6 months (*12*), we next aimed to determine whether CD8^+^CD103^-^CD69^+^CD49a^-^ T_RM_ in the mLN were exhausted effector T cells with a stem-like T_RM_ phenotype. Mice were given IT vaccination with either SSI low dose or BCG::RD1 and mLN of experimental animals harvested across 70, 90, and 120 dpv before processing into single cell suspensions. Exhausted CD8^+^ T cell subsets (*57, 64, 65*) were then quantified using flow cytometry.

After both IT vaccinations we observed cells in the mLN with a CD69^+^CXCR5^+^PD-1^+^ phenotype that are reported to be quiescent stem-like T_RM_ (Fig. 4f) as well as cells with a CD69^-^ CXCR5^+^PD-1^+^ phenotype that are reported to be actively cycling (Fig. 4g). These highly proliferative cells gave rise to reinvigorated cytotoxic CD69^-^CXCR5^-^PD-1^+^ effector T cells (Fig. 4h) and CD69^+^CXCR5^-^PD-1^+^ effector T cells (Fig. 4i). Both subsets of progeny are reported to regain access to circulation, and the latter ultimately may become new T_RM_ of non-lymphoid infected tissues (*57, 64, 65*). Although functional characterisation of these memory T cell subtypes could not be carried out as a part of the present study, data across the 7-week timeline indicated that following vaccination with BCG SSI low dose an average 40% of all phenotypic stem-like T_RM_ cells were proliferative and produced an average amount of ∼13,000 reinvigorated circulatory memory T cells of which ∼5,000 were mildly cytotoxic and ∼8,000 had the capacity to regain residency within the infected lungs. 70-120 days after BCG::RD1 IT vaccination we observed an average of 44% cycling stem-like T_RM_ produced an average amount of ∼130,000 reinvigorated circulatory memory T cells of which ∼80,000 were mildly cytotoxic and ∼50,000 had the capacity to regain lung T_RM_ status. The proliferative potential of the mLN CD69^-^CXCR5^+^PD-1^+^ cells was appreciable, as one cycling cell produced between 50-200 progeny.

In summary, our study demonstrated, for the first time, that mucosal BCG vaccination induced cells with both ex-lung and stem-like T_RM_ phenotypes to persist in the mLN for at least 4 months. Our discovery of persistent cells in the mLN with stem-like T_RM_ phenotypes may have notable implications for future therapeutic interventions for latent TB.

### Migratory distal airway progenitor cells provide paracrine cues for the induction and persistence of CD8^+^ lung T_RM_ cells following mucosal BCG vaccination

We next investigated putative relationships between lung epithelial progenitor cells and CD8^+^ T_RM_ following mucosal BCG vaccination as we hypothesized that distal airway progenitor cells provide paracrine induction cues for lung CD8^+^ T_RM_. The sum of our analyses thus far inferred that induction and maintenance of lung CD8^+^ T_RM_ was linked to the tissue milieu associated with alveolar damage. As described in influenza mouse models, CD8^+^ T_RM_ form in areas of severe alveolar destruction, precisely the areas into which airway progenitor cells home (*21, 22, 28, 34, 66*). Thus, we first looked at spatial and temporal relationships of Krt8^+^ TP and CD8^+^ T_RM_ using imaging and across all vaccination models we visualized evident structural localization, particularly in areas of severe tissue damage (Fig. 5a). Given that BCG::RD1 was associated with the most severe tissue damage, we used vaccination with BCG::RD1 to dissect mechanistic relationships between distal airway epithelial progenitor cells and immune memory.

**Figure 5.**
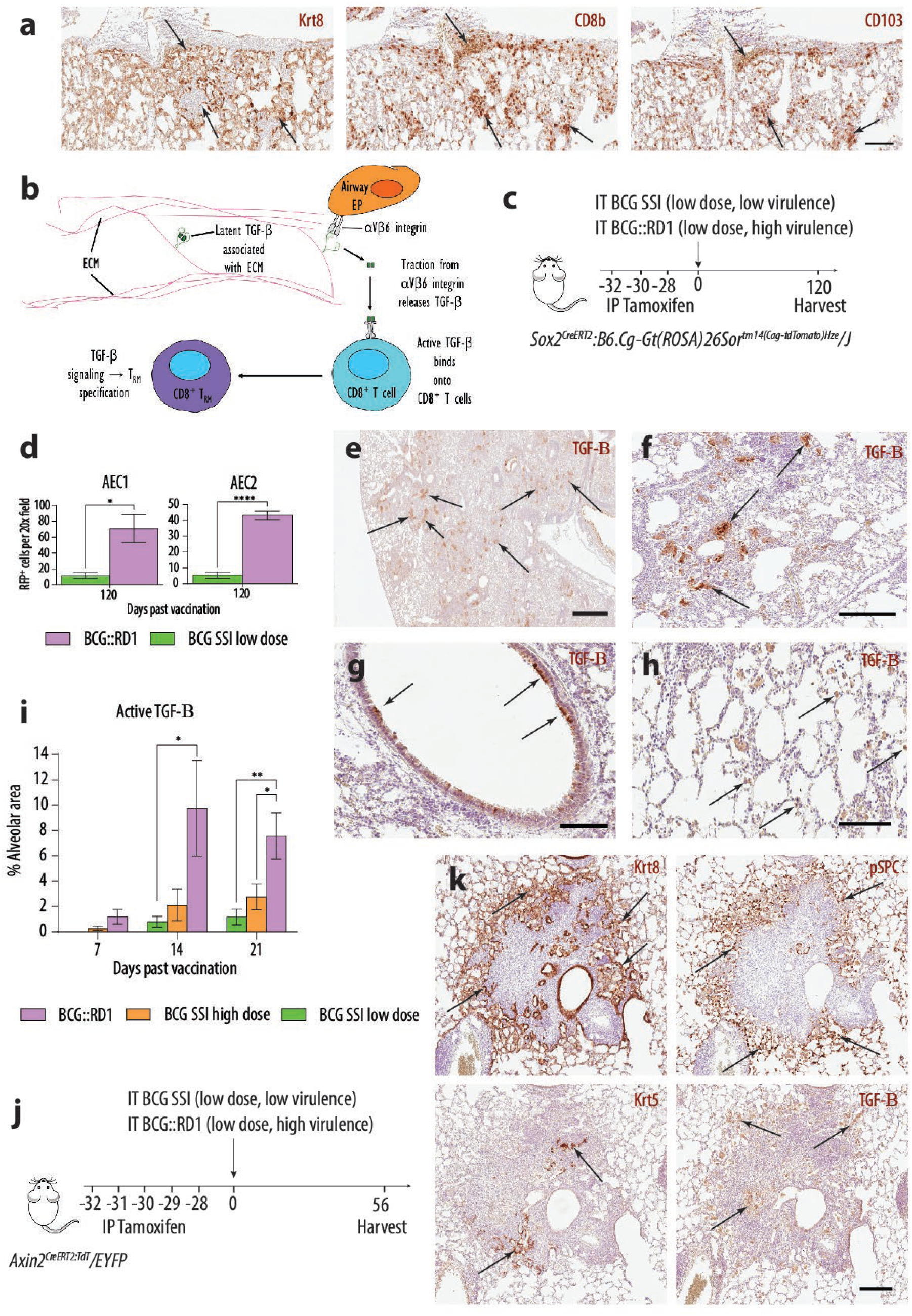
Airway epithelial progenitor migration releases alveolar tissue active TGF-β. **a)** Serial micrographs show localization of Krt8^+^ TP with CD8b^+^CD103^+^ T cells in areas of severe alveolar damage. **b)** Model of interaction between airway epithelial cells and adjacent CD8^+^ T cells. **c)** Experimental design for lineage tracing of *Sox2/Ai14* progeny at 120 dpv and **d)** their quantification. Alveolar interstitial tissue active TGF-β following IT vaccination with BCG::RD1 seen in **e)** distal interstitium (arrows) at 14 days and **f)** areas of severe damage (arrows) at 21 dpv**. g)** Bronchiolar airway expression (arrows) of active TGF-β at 14 days after IT SSI high dose vaccination. **h)** The cytoplasm of alveolar macrophages (arrows) exhibited active TGF-β at 21 days after IT SSI low dose vaccination. **i)** Quantification of extracellular alveolar interstitial active TGF-β following IT BCG vaccinations. Pooled data from two experiments where n=6 for each timepoint. \**P* < 0.05; \*\**P* < 0.01; \*\*\**P* <0.001; \*\*\*\**P* <0.0001; ns: not significant by One-way ANOVA**. j)** Experimental design for lineage tracing of *Axin2^CreERT:TdT^/EYFP* progeny at 120 dpv. **k)** Micrographs of lungs of *Axin2^CreERT:TdT^/EYFP* animals 56 days after mucosal BCG::RD1 vaccination. **a, e-h)** Results are presented from representative images from two independent experiments where n=6 and **k)** n=10. **d)** Pooled data from two experiments where n=8. \**P* < 0.05; \*\**P* < 0.01; \*\*\**P* <0.001; \*\*\*\**P* <0.0001; ns: not significant by unpaired Student’s *t* test. **a)** Magnification, 20x; Scale bar 100µm. **e)** Magnification, 4x; Scale bar 600µm. **f)** Magnification, 11.6x; Scale bar 200µm. **g)** Magnification, 20x; Scale bar 100µm. **h)** Magnification 20x; Scale bar 100 µm. **k)** Magnification 9.6x; Scale bar 150µm.

TGF-β is a key regulator of epithelial cell proliferation and differentiation (*67, 68*) and upregulates the CD8^+^ T_RM_ transcriptome (*69, 70*). A key activator of latent TGF-β in lungs is integrin αVβ6 (*71*) and epithelial progenitors express αVβ6 (*21, 32, 72, 73*) which is used by cells for migration across the extracellular matrix (ECM) (*74–76*). Within the lungs, the latent form of TGF-β is stored in the ECM in high concentrations (*71, 77*) and αVβ6 is known to bind the latent-associated protein (LAP) so that αVβ6-mediated cell traction forces open the LAP to activate TGF-β (*71, 78, 79*). Active TGF-β generated by αVβ6 is utilised by cells in direct contact with αVβ6 expressing cells (*71, 80, 81*). We hypothesized that as airway epithelial progenitor cells migrate into hypoxic alveolar tissue, they activate interstitial latent TGF-β which then binds TGF-β receptors on adjacent CD8^+^ T cells to subsequently induce the CD8^+^ T_RM_ transcriptome (Fig. 5b).

To first test this hypothesis, we quantified lineage traced Sox2^+^ airway progenitors at 120 dpv in regenerated alveolar tissue, using *Sox2^CreERT2^:B6.Cg-Gt(ROSA)26Sor^tm14(CAG-tdTomato)Hze^/J* (aka *Sox2/Ai14*) reporter mice in which progeny of Sox2^+^ airway progenitors were indelibly tagged with red fluorescent protein (Fig. 5c). To ensure there was no tamoxifen persistence, tamoxifen was administered 28 days prior to IT vaccination.

We observed that numbers of lineage traced AEC1 and AEC2 following BCG::RD1 vaccination were 6 and 8-fold higher, respectively, than those observed after BCG SSI low dose. (Fig. 5d); therefore, the number of airway epithelial progenitors in the alveolar tissue was significantly higher following mucosal BCG::RD1 vaccination. We next visualized and quantified extracellular interstitial alveolar tissue active TGF-β from 7-21dpv when airway epithelial progenitors were mobilized and CD8^+^ T_RM_ developed. Results showed that by 7 dpv the BCG::RD1 vaccinated animals had peribronchiolar interstitial active TGF-β expression (data not shown) and by 14 dpv alveolar interstitial expression was evident (Fig. 5e, i). By 21 dpv, alveolar interstitial expression was global, and areas of severe damage had strong expression (Fig. 5f, i).

Animals vaccinated with BCG SSI high dose had sparse peribronchiolar expression of active TGF-β from 14-21 dpv (Fig. 5i) with early, strong extracellular airway expression (Fig. 5g); in animals vaccinated with the low dose of BCG SSI, scarce alveolar macrophages had cytoplasmic expression of active TGF-β (Fig. 5h). Therefore, we revealed that after BCG::RD1 vaccination, active TGF-β was significantly more concentrated in alveolar tissues at the time when airway epithelial progenitors were known to restitute severely damaged epithelium.

We also sought to quantify lineage traced AEP at 120 dpv in regenerated alveolar tissue using *Axin2^CreERT2:TdT^/EYFP* reporter mice to indelibly tag AEP progeny with enhanced yellow fluorescent protein (Fig. 5m). At tissue harvest at 120 dpv, however, all animals had an atypical response to the BCG::RD1 across two independent experiments, despite no underlying genetic abnormality. Numerous lesions of these lungs contained Krt8^+^ TP, AEP and interestingly, Krt5^+^ DASC progeny as well as profuse amounts of alveolar interstitial active TGF-β. Wounds exemplified lung fibrosis due to AEC2 hyperplasia, lack of AEC1 and supraphysiologic levels of TGF-β that likely prevented differentiation of Krt8^+^ TP into AEC1 (*32, 82*) (Fig. 5n).

The observed Krt5^+^ cells in dysplastic lung lesions were primarily interstitial and not adjacent to bronchioles (as described in the murine PR8 influenza model) and we suggest that DASC were activated and mobilized in these animals due to the severity of lung injury. This notion is supported by human explant and autopsy studies of lung tissue following SARS-CoV-2 infection in which non-resolvable chronic lung disease resulted in the recruitment of Krt5^+^ cells adjacent to fibrotic areas of lung long after clearance of infectious virus (*25, 26, 83*). We did not observe the development of dysplasia following any of the other three IT vaccinations with BCG; rather, in the C57BL/6, *Sox2^CreERT2^/Ai14*, and *Krt5^CreERT2^* mice, the epithelial progenitors that responded to injury successfully transitioned from the Sox2^+^ airway and pSPC^+^ alveolar phenotype through the Krt8^+^ TP state and differentiated into AEC1 (evidenced by overall lung restitution). This implied that TGF-β signaling necessary for early upregulation of *Krt8* and induction of the Krt8^+^ TP state was appropriate and that subsequent inactivation of TGF-β promoted terminal differentiation into AEC1 (*82*). In the future, the IT BCG vaccinations employed here may be of use to characterize the pathways of successful reparative processes as opposed to **Figure 5**. Airway epithelial progenitor migration releases alveolar tissue active TGF-β.

### *Itgβ6 knock out* mice develop significantly fewer CD8^+^ T_RM_ following mucosal vaccination with BCG::RD1

To more directly test our theory that epithelial progenitor cell-mediated release of TGF-β regulates the induction of CD8^+^ T_RM_, we used knock-out (KO) mice that lacked a functional αVβ6 integrin (*86, 87*). *Integrin beta-6 (ItgB6) KO* mice were vaccinated with BCG::RD1 and tissues harvested across 7-35 dpv (Fig. 6a). We confirmed levels of injury in the *ItgB6 KO* were comparable to C57BL/6 wild type mice using histology (data not shown) and quantification of Krt8^+^ TP numbers (Fig. 6b). In alveolar tissue, we then compared extracellular interstitial active TGF-β, as *ItgB6 KO* airway progenitor cells would not be able to bind the LAP and activate TGF-β. At 21 dpv (the peak of Krt8^+^ progenitors), we observed that C57BL/6 mice had abundant interstitial active TGF-β, compared to the *ItgB6 KO* animals which had negligible amounts (Fig. 6c, d).

**Figure 6.**
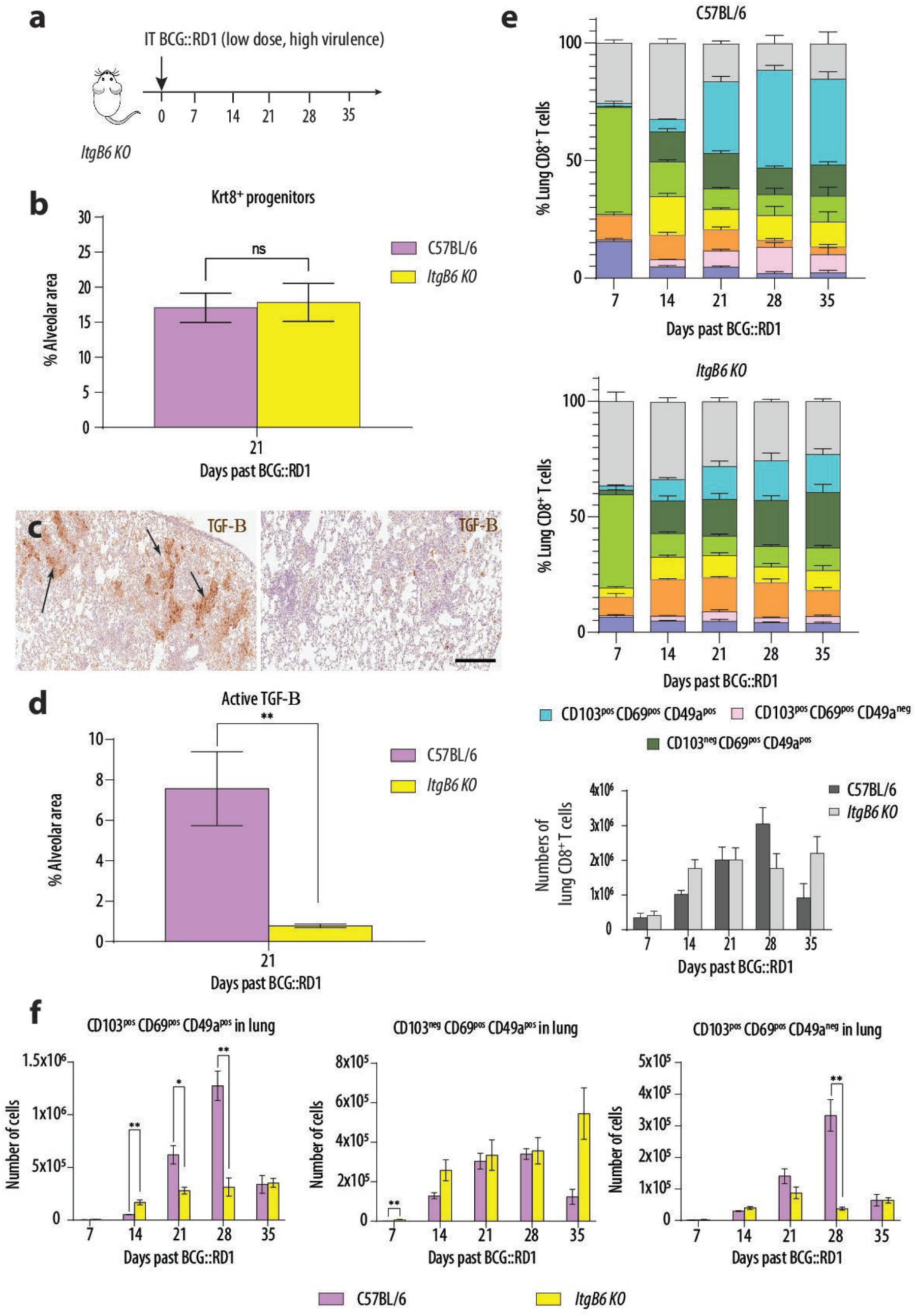
*Itgβ6 KO* mice demonstrate significantly decreased numbers of CD8^+^ T_RM_ following mucosal BCG::RD1 vaccination. **a)** Experimental design for *Itgβ6 KO* experiments. **b)** Quantification of Krt8^+^ TP in *Itgβ6 KO* mice. **c)** Micrographs of lungs at 21 days after BCG::RD1 vaccination and **d)** corelative quantification of extracellular interstitial expression (**c**-arrows) of active TGF-β in alveolar tissue. **e)** Percentages of CD8^+^ T_RM_ subsets following mucosal BCG::RD1 vaccination and numbers of CD8^+^ T cells in the lungs of both strains. **f)** Flow cytometry quantification of numbers of CD8^+^ T_RM_ subsets. Pooled data from two independent experiments where n=6-10 for each timepoint. \**P* < 0.05; \*\**P* < 0.01; ns: not significant by unpaired Student’s *t* test. **c)** Magnification, 10x; Scale bar 200µm.

We next quantified CD8^+^ T_RM_ subsets and their proportions in the lungs of the *ItgB6 KO* animals after BCG::RD1 vaccination across 5 weeks and found that while C57BL/6 mice had a 25% increase in all CD8^+^ T_RM_ between 14-21 dpv, the *IgB6 KO* animals only had an increase of 5% Fig. 6e). This smaller increase was despite the fact that the KO mice had more lung CD8^+^ T cells at 14 dpv and the same number as WT at 21 dpv (Fig. 6e). We also observed that CD8^+^CD103^+^ T_RM_ were specifically significantly decreased in *ItgB6 KO* animals after BCG::RD1 vaccination compared to WT animals (Fig. 6f), consistent with the role of TGF-β in upregulation of CD103 expression in CD8^+^ T cells (*69, 88*).

In conclusion, following mucosal BCG vaccination, wherever Krt8^+^ TP were, CD8^+^CD103^+^ T_RM_ also localized. Active TGF-β was robust and widespread throughout the alveolar interstitium following BCG::RD1 vaccination in which significantly higher numbers of distal airway progenitors were shown. When αVβ6 integrin was knocked out, active TGF-β was absent and CD8^+^CD103^+^ T_RM_ numbers significantly decreased. Taken together, these findings support a model in which lung epithelial progenitor cell-mediated release of TGF-β regulates induction and persistence of lung CD8^+^ T_RM_ cells following mucosal BCG vaccination.

## DISCUSSION

TB is one of the major causes of human death in the world and a new vaccine that induces lung-resident immunity is urgently needed. This is, to our knowledge, the first study to show a mechanistic relationship between lung epithelial progenitor cells and T_RM_ induction/maintenance following mucosal mycobacterial vaccination. Our findings hold several important implications for development of a more effective TB vaccine.

Firstly, mucosal delivery of BCG is known to induce better systemic and mucosal immune responses compared to parenteral BCG (*9, 11, 89, 90*). Our findings further support that inhaled BCG not only led to robust induction of lung CD8^+^ T_RM_ and but also significant populations of cells with an ex-lung T_RM_ and stem-like T_RM_ phenotype that persisted for up to 4 months. Future studies should investigate whether antigen-specific mLN T_RM_ can be harnessed to provide a regional anti-TB arsenal for the lungs.

We also demonstrated that a high dose of BCG SSI delivered to the proximal airways induced 4 times the number of airway CD8^+^ T_RM_ compared to the BCG SSI low dose that affected the distal airways. If the mucosal route is inevitable for a TB vaccine, one engineered with larger particulate matter, delivered to the upper airways (*91*) and ultrafine particles to the distal airways (*92*) could have advantages.

A key focus of TB vaccine improvement is BCG modifications that generate persistent CD8^+^ T cell memory. The BCG::RD1 construct used in our studies has been shown to protect animals from TB challenge following IT administration; however, when bacilli infect AEC2, ESAT-6 causes cell necrosis to facilitate spread into the interstitium (*93*). We confirmed that RD1virulence factors had a direct influence on the severity and extended time of AEC losses, inflammation, CD8^+^ T cell responses and ultimately, tissue regeneration. Importantly, epitopes of a pulmonary mucosal vaccine should be proven not to facilitate BCG adherence and internalisation into AEP. These crucial progenitor cells that function as AEC2 (*23*) may be frontline cells for BCG invasion and it is known that infection can impede epithelial stem/progenitor cell function, leading to decreased (*94*), faulty (*95*), or incorrectly timed production of progeny (*96*), and subsequent development of lung disease (*97*).

At the time of our experimental design, the standard BCG dose for mouse work was 500,000 CFU since the World Health Organisation recommends use of high dose vaccination for children (*98*). Mouse studies have shown, however, that higher doses of BCG does not correlate with better protection (*99, 100*); although we administered BCG::RD1 at 100,000 CFU, BCG SSI administered at 3,000 CFU has demonstrated protective efficacy in recent mouse studies (*101*). As BCG::RD1 stimulated a robust local and regional immune memory, we will endeavour to continue studies with the 3,000 CFU dose to ameliorate severity of lung pathology and study protection against *Mtb* challenge. Also, mycobacteria are slow growing and the time to generate immunology data is long; therefore, most studies have been limited to 2-3 months. Future vaccine efficacy studies should extend to 6 months or a year (*9*) before challenge with *Mtb*, as a bi-annual or annual dose of vaccine would be practical for humans.

We demonstrated the importance of the early resolution of inflammation and downregulation of TGF-β as this corresponds to the turning point for lung regeneration (*102*). Clearly, a vaccine that leads to excessive inflammation, while it may induce more CD8^+^ T_RM_, is detrimental to the lungs. We learned that excessive AEC losses mobilised airway epithelial progenitors and this was found directly related to TGF-β activation in bronchiolar epithelium and alveolar interstitium. Although this may have resulted in more CD8^+^ T_RM_, increased levels or time of activated TGF-β was also shown to harm the lungs. Therefore, an optimal pulmonary mucosal vaccine should enhance local epithelial progenitor responses and keep interstitial active TGF-β levels low so as not to tip the balance between CD8^+^ T_RM_ induction and tissue integrity (*68, 71, 72, 102–105*).

Although there remains optimism in the field that numbers of vaccine induced CD8^+^ T_RM_ against *Mtb* could be increased and made to persist in the lungs (*106*), our results add further evidence that CD8^+^ T_RM_ have a distinctive *lack* of endurance in the lungs compared to other tissues (*21, 49, 53*) which is thought to be organ specific (*21, 49, 53*). A future research direction for a new pulmonary mucosal TB vaccine may be to develop one that minimises alveolar harm (and active TGF-β), to generate lung CD103^-^CD69^+^CD49a^+^ T_RM_ which are known to specifically egress to the mLN. Since mLN CD103^-^CD69^+^CD49a^+^ CD8^+^ T_RM_ may have the capacity to migrate back to the lungs and orchestrate a timely and efficient innate response (*54–56*), it could be beneficial to study whether those generated from the mucosal vaccination persist in the mLN for six months to a year with the continued capacity to protect the lungs against *Mtb* challenge in the interim.

### Limitations of results and interpretations

Our decision not to use intravascular labelling to discriminate between CD8^+^ T cells in the lung circulation as opposed to the parenchyma was since across our vaccinations, we observed vascular leakage (extensive in many animals) where exudation of the antibodies into the tissues would have compromised interpretation of the staining. This decision was supported by, previous studies that have shown that CD69^+^ and CD103^+^ T cells were found only in lung non-lymphoid parenchyma and were protected from intravascular labelling (*107, 108*).

Limitations to our experiments will require new tools and additional work to address. A major limitation of the study is that Sox2 is ubiquitously expressed in all airway epithelial cells and precluded us from discerning the precise stem cell origin of Sox2^+^ progenitor cells. (*16, 22, 30, 31, 109–111*). Future studies should employ parabiosis to prove that the CD8^+^ T_RM_ harboured in the mLN after IT vaccination have bona fide mLN residency and tetramer labelling should be used in all further studies to assure mLN CD8^+^ T_RM_ are vaccine specific.

## MATERIALS AND METHODS

### Study Design

Mucosal BCG delivery enhances CD8^+^ T_RM_ generation protection against *Mtb* challenge (*11, 112*), thus we sought to delineate mechanistic relationships between lung stem cell progeny and CD8^+^ T_RM_ for possible translation into the safe improvement of TB vaccine strategies.

In controlled lab experiments, C57BL/6 mice were IT vaccinated with various doses and strains of *M. bovis* BCG across a spectrum of virulence. IHC, imaging, and flow cytometry were used to prospectively characterise the timeline of stem cell activation and CD8^+^ T_RM_ differentiation. Numbers of migratory airway progenitors that restituted alveolar tissue were quantified using lineage tracing and correlations between activated TGF-β, epithelial progenitors and CD8^+^

T_RM_, was determined using IHC and flow cytometry where analyses defined their structural colocalization; *ItgB6 KO* mice were used to inhibit airway epithelial progenitor activation of TGF-β. Samples were obtained from 4-10 animals for each timepoint investigated; experiments were replicated 2 times. This study (A2837) was approved by the James Cook University Animal Ethics Committee.

### Animals

Female mice were 6–8 weeks old at the time of vaccination and maintained in a biosafety level 2 facility under specific pathogen free conditions. C57BL/6 mice sourced from the ARC (Western Australia), *Krt5^CreERT2^* mice provided by Dr Felicity Davis (University of Queensland), *Axin2^CreERT2:TdT^/EYFP* mice provided by Dr Edward Morrisey (University of Pennsylvania), *Sox2^CreERT2^* (Jackson Laboratories), *Itgβ6* knockout mice (provided by Dr Dean Sheppard, University of California San Francisco), *B6.Cg-Gt(ROSA)26Sor^tm14(CAG-tdTomato)Hze^/J* (also known as *Ai14*) mice (provided by Dr Scott Mueller, University of Melbourne), *Gt(ROSA)26Sor^tm4(ACTB-tdTomato,-EGFP)Luo^*/J (also known as *mTmG*) mice originally from Jackson Laboratories (*37*) with substrain donated by Dr Scott Mueller (University of Melbourne) were all genotyped by Genetic Research Services (University of Queensland), bred and maintained in the animal facilities at James Cook University, Australia.

### Bacteria

BCG SSI (ATCC no. 35733) and BCG::RD1 (provided by Dr Roland Brosch, Institut Pasteur) were grown in Middlebrook 7H9 broth (BD Biosciences) supplemented with 0.2% glycerol, 0.05% Tween 80, and 10% ADC enrichment (BD Biosciences) as previously described (*13*).

### Mucosal Vaccination

Mice were immunized by IT instillation of 1 x 10^5^ or 5×10^5^ colony forming units (CFU) of BCG diluted in sterile PBS as described previously (*11*). Animals were monitored on a weekly basis, with determination of weight and scoring of body condition using the mouse BCS scale. At the doses utilized in this study, none of the animals met institutional animal care and use committee approved endpoints and they were analysed at the planned time-points.

### Lung Inflation

After euthanasia by CO_2_ inhalation, animals were exsanguinated; the thorax was opened to expose the lungs. An 18-gauge stub needle tip was inserted into the trachea and secured. The needle was connected to a reservoir with 0.5% zinc acetate and zinc chloride in 0.5% tris-calcium acetate buffer (*113*) (ZBF; BD Biosciences). Lungs were inflated with ZBF at a fluid pressure of 25cm before removal *en bloc* and incubated in 30ml ZBF at 25°C for 72 hr prior to 24-hr incubation at 25°C in 50% ethanol.

### Immunohistochemistry

Fixed tissues were dehydrated with an automated tissue processor (Leica Biosystems) and embedded in paraffin blocks. 4µm sections were cut and collected on Polysine adhesion slides (ProSciTech), dried at 25°C overnight prior to deparaffinization using standard protocols.

H&E staining was performed using an automated processor (Leica Biosystems). Sections were washed each time with Tris-buffered saline (TBS) and blocked with 10% species-specific serum to secondary antibodies at 25°C for 2 hr. Primary antibodies were incubated overnight at 4°C in optimized dilutions (Supp. Table 1). Tissues were washed before incubation with peroxidase inhibitor (Thermo Fisher Scientific) for 30 min at 25°C. For avidin biotin complex (ABC) IHC, endogenous avidin binding activity was quenched with avidin and biotin blocking buffers (Abcam) for 15 min at 25°C, and tissue washed before incubation with secondary antibodies for 1 hr at 25°C in optimized dilutions (Supp. Table 1).

For ABC IHC, tissue was washed before incubation with 45µl of Vectastain ABC kit reagents (Vector Laboratories) in 10ml TBS for 30 min at 25°C. Tissue was washed before incubation with DAB chromogen (Abcam) for 3-5 min, washed in distilled water and counterstained with hematoxylin (Sigma-Aldrich). Tissues were cleared through ethanol and xylene changes using standard protocols, cover slipped with Surgipath mounting media (Leica) and #1 coverslips (Leica).

### Microscopy

Images were acquired using an Aperio CS2 Digital Slide Scanner (Leica Biosystems) and analyzed using Aperio ImageScope v12.4.3.5008 software (Leica Biosystems). Quantification of Krt8^+^ TP, active TGF-β and Sox2 lineage traced RFP+ cells was undertaken using ImageJ software (ImageJ).

### Lung and Mediastinal Lymph Node Dissociation

Lung and spleen dissociation kits (Miltenyi Biotec) were used to dissociate lung and mLN into single cell suspensions, respectively, according to manufacturer’s instructions. After the final dissociation step, supernatant was aspirated before addition of 3ml RBC lysis buffer (Thermo Fisher Scientific), resuspension and incubation for 2 minutes. 10ml FACS buffer was added to stop reaction prior to centrifugation at 1500 rpm for 5 minutes at 25°C. Supernatant was discarded and cells resuspended with FACS buffer to 500µl.

### Flow Cytometry and Data Analyses

150µl of dissociated cells were pipetted into 96 round bottom well plate and spun at 1500 rpm at 4°C for 5 minutes. Supernatant was discarded and cells resuspended in 50µl Live/Dead cell stain 1:1000 (APC-Cy7 channel) (BD Biosciences) for incubation at 25°C for 10 minutes in the dark. Samples were washed with FACS buffer, spun at 1500 rpm for 5 minutes at 4°C.

Supernatant was discarded and samples resuspended in 50µl antibody master mix consisting of FACS buffer, and antibodies at optimized concentrations (Supp. Table 1). Samples were incubated for 30 minutes on ice in the dark, washed twice with FACS buffer, spun at 1500 rpm at 4°C for 5 minutes and supernatant discarded. 150µl Sphero Blank Calibration Particles (BD Biosciences) at 1:75 was added to resuspend samples. Flow cytometry analyses were performed on a LSR Fortessa cytometer using FACSDiva software (BD Biosciences). The data were analysed by FlowJo software (Treestar).

### Lineage Tracing

A single dose of 0.125mg/g body weight tamoxifen (Sigma-Aldrich) was dissolved in 50µl corn oil (Sigma-Aldrich) and administered to *Krt5^CreERT2^/mTmG* mice intraperitoneally at day 14 after IT vaccination with BCG::RD1. Three doses of 0.25mg/g tamoxifen was administered every second day to *Sox2^CreERT2^/Ai14* mice and every day for 5 days to *Axin2^CreERT2:TdT^/EYFP* mice 28 days prior to IT vaccinations. Tamoxifen-independent recombination in *Krt5^CreERT2^/mTmG* mice was 0% with no systemic effects in floxed littermate controls. In *Sox2^CreERT2^/Ai14* mice tamoxifen independent recombination was less than 1%, while floxed controls were negative for systemic effects. *Axin2^CreERT2:TdT^/EYFP* were unable to be analysed for independent recombination due to excessive lung pathology.

### Statistical Analysis

Statistical analysis and graphs were generated using Prism version 9.5.1 (GraphPad). Two parametric group analyses were carried out using Student’s t test, followed by Holm-Šídák multiple comparison tests. Analyses of variance between independent groups were carried out using One-way ANOVA by Tukey’s multiple comparison tests. *P* < 0.05 was considered significant.

## Acknowledgments

We thank M. Stafford and J. Whan for technical assistance; F. Davis, E. Morrisey, D. Sheppard, S. Mueller for generously sharing mouse lines; University of Queensland Genetic Research Services for genotyping animals and Developmental Studies Hybridoma Bank for reagents.

## Funding

James Cook University Prestige Scholarship (JAB)

National Health and Medical Research Council (NHMRC) Investigator grant GNT2008715 (AK)

National Health and Medical Research Council (NHMRC) Ideas grant GNT2001262 (AK)

### Author contributions

J.A.B., P.R.G., D. L. D. and A. K. conceived, designed, and analysed experiments and wrote the manuscript. J.S. performed IT vaccinations; R.R. performed IP tamoxifen injections. J.A.B. performed all other animal work, IHC, flow cytometry and imaging.

### Data and materials availability

All data are available in the main text or the supplementary materials.

## Supplementary Materials

**Supplementary Figure 1.**
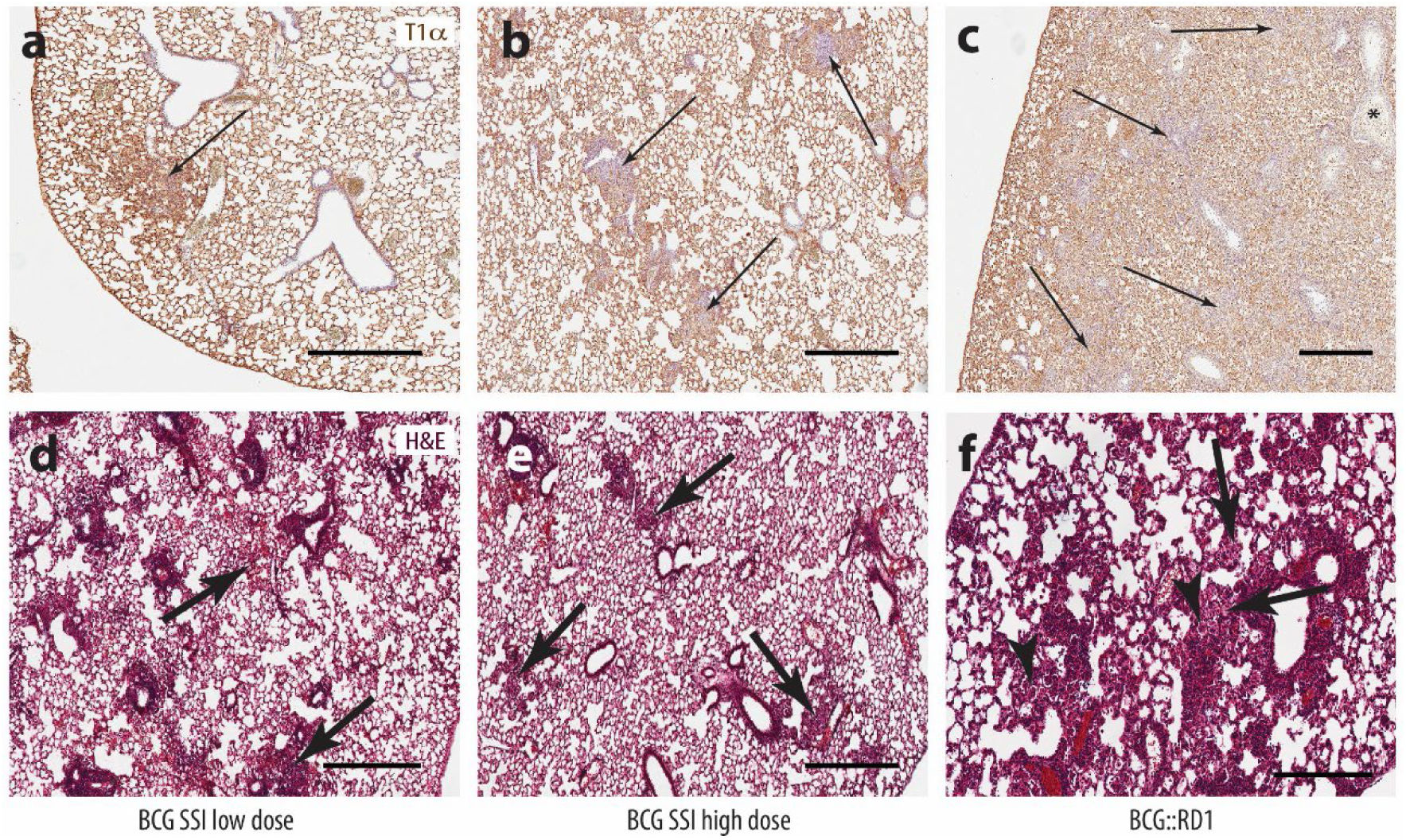
a-c) Lesions with loss of T1α AEC1 marker staining (blue areas indicated by arrows) due to vaccine pathology. **c)** Necrotic AEC debris within bronchiole lumen (indicated by asterisk) due to BCG::RD1 vaccination. **d)** H&E staining shows focal areas of hemorrhage (arrows) due to the SSI low dose vaccination, **e)** interstitial giant cells (arrows) due to the SSI high dose vaccination **f)** alveolar giant cells (arrowheads) and foamy cells (arrows) due to the BCG::RD1 vaccination. Results are presented from representative images from 6 mice per group and pooled data means ± SEM from two independent experiments. **a-c)** Magnification, 6x; Scale bar 400 µm. **d)** Magnification, 5x; Scale bar 500 µm. **e)** Magnification, 4.8x; Scale bar 500 µm. **f)** Magnification, 10x; Scale bar 300 µm.

**Supplementary Table 1.** List of antibodies used for IHC, IF and flow cytometry.

| Primary Antibody | Species raised | Manufacturer/ Concentration | Working dilution | Secondary Antibody | Species raised | Manufacturer/ Concentration | Working dilution |
| --- | --- | --- | --- | --- | --- | --- | --- |
| T1α<br>8.1.1-s | Hamster serum | DSHB<br>15ug/ml | 2ug/ml | Biot-Hamster IgG<br>ab6782 | Rabbit | abcam<br>1.5mg/ml | 1:1000<br>w/2% NRS |
| Pro-SPC<br>ab90716 | Rabbit IgG | abcam<br>0.7mg/ml | 1:1000 | HRP-Rabbit IgG<br>ab97057 | Goat | abcam<br>0.5mg/ml | 1:50 |
| Krt5<br>Poly9059 | Chicken IgY | BioLegend<br>0.5 mg/mL | 1:200 | Biot-Chicken IgY<br>Ab6876 | Goat | abcam<br>2.0mg/ml | 1:800 |
| Sox2<br>EPR3131 | Rabbit IgG | abcam<br>0.214mg/ml | 1:200 | Biot-Rabbit IgG<br>ab6720 | Goat | abcam<br>2.0mg/ml | 1:1000 |
| Krt8<br>EP1628Y | Rabbit IgG | abcam<br>0.033mg/ml | 1:1000 | HRP-Rabbit IgG<br>ab97057 | Goat | abcam<br>0.5mg/ml | 1:50 |
| CD3e<br>EPR22667-12 | Rabbit IgG | abcam<br>0.673mg/ml | 1:100 | HRP-Rabbit IgG<br>ab97057 | Goat | abcam<br>0.5mg/ml | 1:50 |
| CD8b<br>EPR22331-54 | Rabbit IgG | abcam<br>0.63mg/ml | 1:500 | HRP-Rabbit IgG<br>ab97057 | Goat | abcam<br>0.5mg/ml | 1:50 |
| CD103<br>EPR22590-27 | Rabbit IgG | abcam<br>0.516mg/ml | 1:200 | HRP-Rabbit IgG<br>ab97057 | Goat | abcam<br>0.5mg/ml | 1:50 |
| TGF-beta<br>1/1.2<br>AF-101-NA | Chicken IgY | R&D Systems<br>0.2mg/ml | 1:100 | Biot-Chicken IgY<br>Ab6876 | Goat | abcam<br>2.0mg/ml | 1:800 |
| RFP<br>600-401-379 | Rabbit IgG | Rockland<br>1.06mg/ml | 1:1000 | HRP-Rabbit IgG<br>ab97057 | Goat | abcam<br>0.5mg/ml | 1:100 |
| GFP | Goat IgG | Aves Labs<br>10mg/ml | 1:500 | HRP-Goat IgG<br>A15999 | Donkey | Thermo Fisher<br>1 mg/ml | 1:500 |
| CD8b<br>EPR22331-54 | Rabbit IgG | abcam<br>0.63mg/ml | 1:500 | AF647-Rabbit IgG<br>AB_2536183 | Donkey | Invitrogen<br>2 mg/ml | 1:1000 |
| CD103<br>EPR22590-27 | Rabbit IgG | abcam<br>0.516mg/ml | 1:200 | AF488-Rabbit IgG<br>AB_2536183 | Goat | Invitrogen<br>2 mg/ml | 1:1000 |
| Krt8<br>EP1628Y | Rabbit IgG | abcam<br>0.033mg/ml | 1:1000 |  |  |  |  |
| TGF-beta<br>1/1.2<br>AF-101-NA | Chicken IgY | R&D Systems<br>0.2mg/ml | 1:100 |  |  |  |  |
| CD49a/BV421<br>Ha31/8 | Rat IgG | BD Biosciences<br>0.2mg/ml | 1:200 |  |  |  |  |
| CD8a/FITC<br>53-6.7 | Rat IgG | BD Biosciences<br>0.5mg/ml | 1:200 |  |  |  |  |
| CD69/PE<br>H1.2F3 | Rat IgG | BD Biosciences<br>0.2mg/ml | 1:100 |  |  |  |  |
| CD103/AF647<br>2E7 | Rat IgG | BD Biosciences<br>0.2mg/ml | 1:200 |  |  |  |  |
| CD16/32<br>2.4G2 | Rat IgG | BD Biosciences<br>0.5mg/ml | 1:500 |  |  |  |  |
| CD185/BV421<br>2G8 | Rat IgG | BD Biosciences<br>0.2mg/ml | 1:50 |  |  |  |  |
| CD279/APC<br>Lγ-51 |  | BD Biosciences<br>0.2mg/ml | 1:200 |  |  |  |  |

